# Human Substantia Nigra Neurons Encode Reward Expectations

**DOI:** 10.1101/2024.05.10.593406

**Authors:** Zarghona Imtiaz, Ayaka Kato, Brian H. Kopell, Salman E. Qasim, Arianna Neal Davis, Lizbeth Nunez Martinez, Matt Heflin, Kaustubh Kulkarni, Amr Morsi, Xiaosi Gu, Ignacio Saez

## Abstract

Dopamine (DA) signals originating from substantia nigra (SN) neurons are centrally involved in the regulation of motor and reward processing. DA signals behaviorally relevant events where reward outcomes differ from expectations (reward prediction errors, RPEs). RPEs play a crucial role in learning optimal courses of action and in determining response vigor when an agent expects rewards. Nevertheless, how reward expectations, crucial for RPE calculations, are conveyed to and represented in the dopaminergic system is not fully understood, especially in the human brain where the activity of DA neurons is difficult to study. One possibility, suggested by evidence from animal models, is that DA neurons explicitly encode reward expectations. Alternatively, they may receive RPE information directly from upstream brain regions. To address whether SN neuron activity directly reflects reward expectation information, we directly examined the encoding of reward expectation signals in human putative DA neurons by performing single-unit recordings from the SN of patients undergoing neurosurgery. Patients played a two-armed bandit decision-making task in which they attempted to maximize reward. We show that neuronal firing rates (FR) of putative DA neurons during the reward expectation period explicitly encode reward expectations. First, activity in these neurons was modulated by previous trial outcomes, such that FR were greater after positive outcomes than after neutral or negative outcome trials. Second, this increase in FR was associated with shorter reaction times, consistent with an invigorating effect of DA neuron activity during expectation. These results suggest that human DA neurons explicitly encode reward expectations, providing a neurophysiological substrate for a signal critical for reward learning.

## Introduction

The dopaminergic system is crucial for reward-based learning and motivational vigor. Activity of individual dopamine (DA) neurons in the ventral tegmental area (VTA) and substantia nigra (SN) encodes reward prediction errors (RPE)^1,2^. RPEs reflect the difference between actual and expected reward and constitute an efficient and robust signal underlying behavioral adaptation in uncertain environments. Abundant evidence across species^1,3–12^ supports the notion that dopaminergic activity, including the spiking rates of individual DA neurons, encode RPEs. However, less is known about how the reward expectations that subserve RPE calculations are encoded in the dopaminergic system. Studies in animal models have found electrophysiological correlates of reward expectation in various brain regions, including the caudate nucleus^13^, striatum^9^, orbitofrontal cortex (OFC)^14^, and globus pallidus externus (GPe)^15^; DA neurons in the rodent VTA then combine information about reward outcomes and expectations into a largely homogeneous RPE signal which is broadcast to downstream targets^12,16^. Conversely, DA neuron activity during the expectation period correlates with the expected value of the upcoming action^17,18^, raising the possibility that reward expectations, an important component in the calculation of RPEs, are explicitly and separately encoded in DA neuron activity. Importantly, competing models of dopaminergic reward-related activation make different predictions about the representation of reward expectation signals in DA neurons: if integration happens at the level of DA neurons, both expectation and reward outcome signals should be reflected in DA neuron activity; conversely, if these are combined in upstream regions, they may not be separable.

This knowledge gap of how DA neurons integrate information about reward expectation and reward into RPEs is particularly salient in the human literature, due to difficulties of recording activity from deep DA nuclei in living humans. Previous studies have leveraged neurosurgical interventions to examine human dopaminergic activity using microelectrode recordings to record single unit activity from the SN^6,7^, or voltammetry approaches to record dopamine transients in the striatum^19–21^. These studies have demonstrated that post-reward activity in the human dopaminergic system reflects reward signals, including unexpected rewards and RPEs. However, whether and how reward expectation signals are encoded in the human dopaminergic system remains unknown.

Here, we sought to examine whether reward expectation is explicitly encoded in the activity of individual human SN neurons. We recorded single unit activity (SUA) intraoperatively from the SN of patients undergoing deep brain stimulation (DBS) surgery for Parkinson’s disease (PD) while the patient played a two-armed bandit task in which they chose between two slot machines with different outcome probabilities that reversed at certain points in the task. We focused our analyses on pre-outcome activity during the reward expectation period, after choice but before reward reveal, and used computational modeling of patients’ behavior to gain insight into patients’ behavioral strategies. Our computational modeling results indicate that patients followed behavioral strategies in which choices were strongly determined by the outcome of the preceding trial. Accordingly, the outcome of the previous trial impacted neural activity: the FR of putative DA SN neurons was higher following positive (+$10) compared to neutral ($0) and negative (-$10) outcomes. Furthermore, previous outcomes impacted patients’ reaction times (RTs), with faster responses following positive compared to negative/neutral outcomes. Together, these results suggest that the baseline FR of human DA neurons reflects a reward expectation signal, and that increased DA tone is associated with an increase in motivational response vigor. This novel evidence supports the idea that human SN neurons explicitly and separably encode reward expectation information in the human dopaminergic system, and that this dopaminergic activation may mediate a relationship between reward expectations and response vigor.

## Results

We carried out SUA recordings during two-armed bandit play in 13 sessions from Parkinson’s disease patients (n=11; 7 male, 4 female, average age = 65.71 ± 7.69 years) who were undergoing DBS lead implantation in the subthalamic nucleus (STN). The microelectrode tip was placed in the SN, past the ventral border of the STN (see Figure 1A and Methods) to record activity of SN neurons. We carried out 8 recordings from the left side of the brain and 5 recordings from the right side of the brain, with bilateral recordings in two patients (Supplemental Table 1). We recorded from a total of 27 neurons, for an average of 2.077 ± 0.277 neurons per patient. To determine the putative identity of individual neurons, we used two different criteria, FR, and spike width. We classified neurons with a FR below 15 Hz and a waveform width >0.8 ms as putative DA neurons and those with greater than 15 Hz FR and/or a waveform width <0.8 ms as putative GABAergic interneurons^22^. Out of our 27 neurons, 22 were classified as putative DA neurons (average FR = 4.883 ± 4.657 Hz) and 5 neurons were categorized as GABAergic (average FR= 33.414 ± 10.607 Hz).

**Figure 1.**
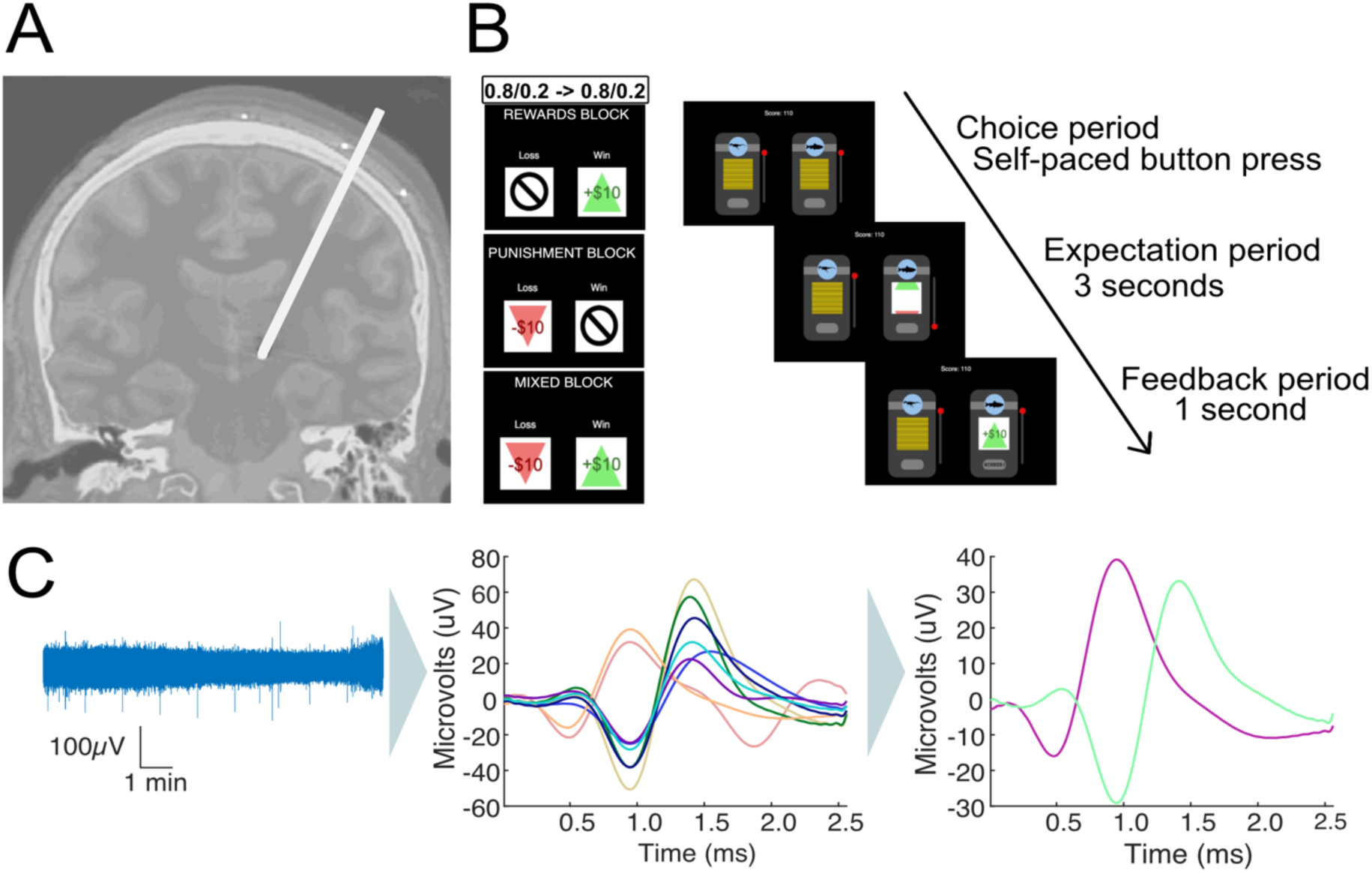
Invasive microelectrode recordings and slot machine task. **(A)** Electrode placement during DBS surgery for Parkinson’s Disease patients. Microelectrodes were parked past the ventral border of the subthalamic nucleus, and single-unit recordings taken from the substantia nigra. **(B)** Two-armed bandit slot machine behavioral task. Patients choose between two slot machines with 80%/20% and 20%/80% probability of a better/worse outcome. Patients play 3x35-trial blocks in pseudo-randomized order, with either +$10/0 (reward block), +$10/-$10 (mixed block) or -$10/$0 (punishment block) outcomes. The contingencies switched twice between slot machines during each block. The patients are shown the slot machines until they make a choice. After a machine is chosen, it spins for 3 seconds until landing on the outcome, which stays on the screen for 1second before the slot machines reset and a new trial starts. **(C)** Microelectrode data (left) is high pass filtered, spike sorted to identify individual units (middle) and manually processed to discard or merge clusters into the final units per recording (right).

Patients played a multi-round two-armed bandit task in which they chose between two slot machines with probabilistic reward outcomes (Figure 1B) with the goal of maximizing their overall reward. The task had three blocks with 35 trials per block: a reward block (+$10/$0 outcomes), a punishment block (-$10/$0 outcomes), and a mixed block (+$10/-$10 outcomes). Block order was randomized across patients. One of the machines had a high probability (80%) of resulting in the better outcome, and the other had a low probability (20%) (Figure 1B). The initial location (left/right) of the high reward machine was randomized, and reward contingencies were reversed twice throughout the task, between trials 12/13 and trials 24/25.

### Computational model reveals patients’ choices were dependent on the outcome of previous trials

We started by characterizing patient behavior using model-free metrics: number of wins and losses per block, as well as the percentage of trials in which the patient picked the machine with the higher reward probability (correct choices). Patients adjusted their behavior after contingency reversals and picked the new machine with the higher reward rate (Figure 2A-C, reward = 67.91±12.29%, mixed = 67.03±12.0%, punishment = 56.57±16.0%). Overall, patients performed better than chance (Figure 2D, p = 3.6583e-11, binomial test), indicating that they understood the game. There was no difference in the proportion of correct choices between blocks (Figure 2E, all p-value >0.05, paired t-test).

**Figure 2.**
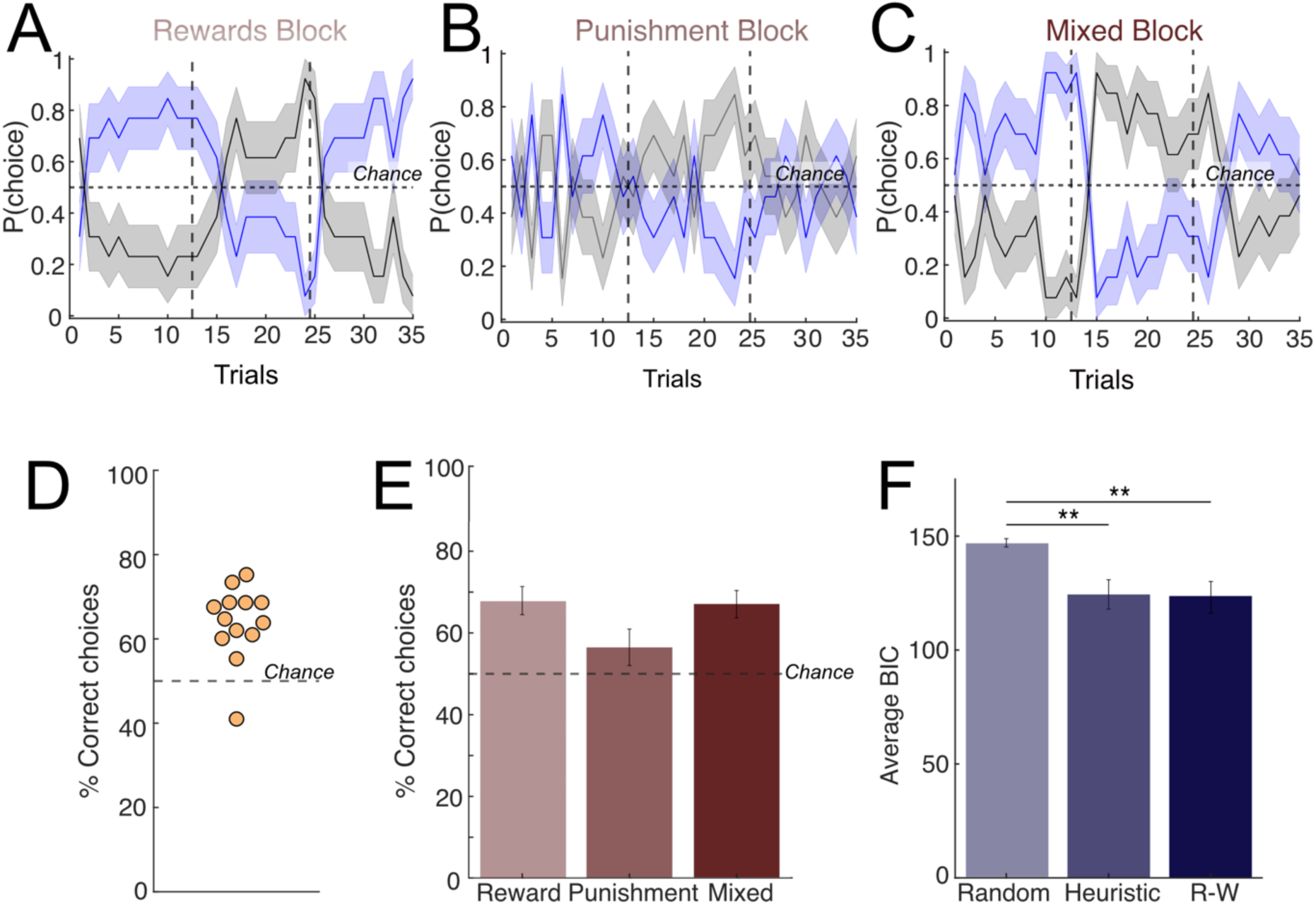
Patient behavior and computational modeling. Average patient behavior across the **(A** rewards, **(B)** punishment, and **(C)** mixed blocks. Patients played 3 blocks corresponding to reward/punishment/mixed conditions in pseudo-randomized order. Within each block, win/loss probabilities (80%/20%) were switched twice (dashed vertical lines). The horizontal solid line represents chance (50%). **(D-E)** Percent of correct choices in which the patient picked the machine with more frequent better outcomes, per patient, across all blocks **(D)** and within each block **(E)**. Dashed line represents chance (50%). The majority of patients selected the correct machine over 50% of the time (n=10/11 patients). **(F)** Computational modeling of behavior comparing the three models considered: a random choice model (Random), a Heuristic win-stay/lose-switch model (Heuristic) and a Rescorla-Wagner model (Rescorla Wagner, RW). The RW produced the best fit to patient behavior (lowest average BIC score) but there was no significant difference between the average BIC scores of the Heuristic and RW models (paired t-test, p- value = 0.7225). Both Heuristic and RW models had a significantly lower BIC score than the random model (paired t-test; heuristic p-value = 0.0046, Rescorla-Wagner p-value = 0.0049).

Next, we sought to further understand the behavioral strategies used by patients through computational modeling of behavior. We fit patient behavior to three different model alternatives: a random model in which patients choose randomly between the two options, a heuristic win-stay/lose-shift model, and a Rescorla-Wagner (RW) model commonly used in RL^23–25^. In the heuristic model, actions are determined by the previous trial outcome (either a ‘good’ or a ‘bad’ outcome, which varied depending on the block identity), using a win-stay/lose-shift rule. Under the RW model, the agent learns from the previous outcomes to determine the values of each slot machine and act accordingly. The RW model produced the lowest average BIC score when fitting the patients’ behavior overall, closely followed by the heuristic win-stay/lose-shift model (Figure 2F; Supplemental Figure 1 for the result for parameter recovery). The estimated learning rate for the RW model for most patients was 1 (mean=0.7390, std=0.4110), indicating that patient choices under this model were heavily dependent on the previous trial. Overall, the computational modeling results indicate that either a RW model with a high learning rate or a heuristic win-stay/lose-switch model are capable of capturing the patients’ behavior. Under either of these strategies, choices are strongly dependent on the outcome of the previous trial.

### Firing rate during expectation is modulated by prior outcome

Next, we sought to examine whether individual SN neurons encode expectation information. We analyzed the FR of putative DA neurons (n=22) during the reward expectation period, after patient choice but before outcome reveal (3s; Figure 1B). Since our modeling results indicated that patients’ choices were strongly determined by prior trial outcomes (under either the RW or heuristic models; Figure 2), we hypothesized that neural signals reflecting reward expectation during this period should depend on previous trial outcome. Because OR time limitations meant that within-block analyses would have resulted in small numbers of trials per block and low statistical power, instead we opted to group the neural activity by previous trial outcome (+10, 0, or -10) independent of block. We observed that neuronal FR during the expectation period was dependent on prior trial outcome, with neurons showing higher FRs in trials following positive trials (+$10) than in negative trials (-$10; two representative examples shown in Figure 3). Some neurons also showed a difference in FR between neutral ($0) and negative (-$10) outcomes, reflecting a gradual increase in expectation period FRs going from negative to neutral to positive outcomes (Figure 3B). To rule out potential confounding factors, we used a generalized linear model (GLM) approach^26^ that included prior motor action (left or right), prior trial reward identity (win or loss), prior trial reward outcome (+10, 0, or -10), and prior trial RPE on the average FR for each neuron as regressors. Only previous reward outcome showed a significant effect on FR during the expectation period (p<10^-4^; all others p>0.05). To examine whether these effects were specific to the expectation period, or whether they reflect sustained FR modulation across trials, we run a similar GLM during the choice period (after slot machine presentation, but before choice). We found no effect of previous trial outcome on FR during choice period (p>0.05), indicating that the sustained FR modulation is specific to the expectation epoch. Finally, we asked whether neuron activity reflected motor activity by examining the average FR during the decision period but observed no movement related modulation at button press (Supplemental Figure 4).

**Figure 3.**
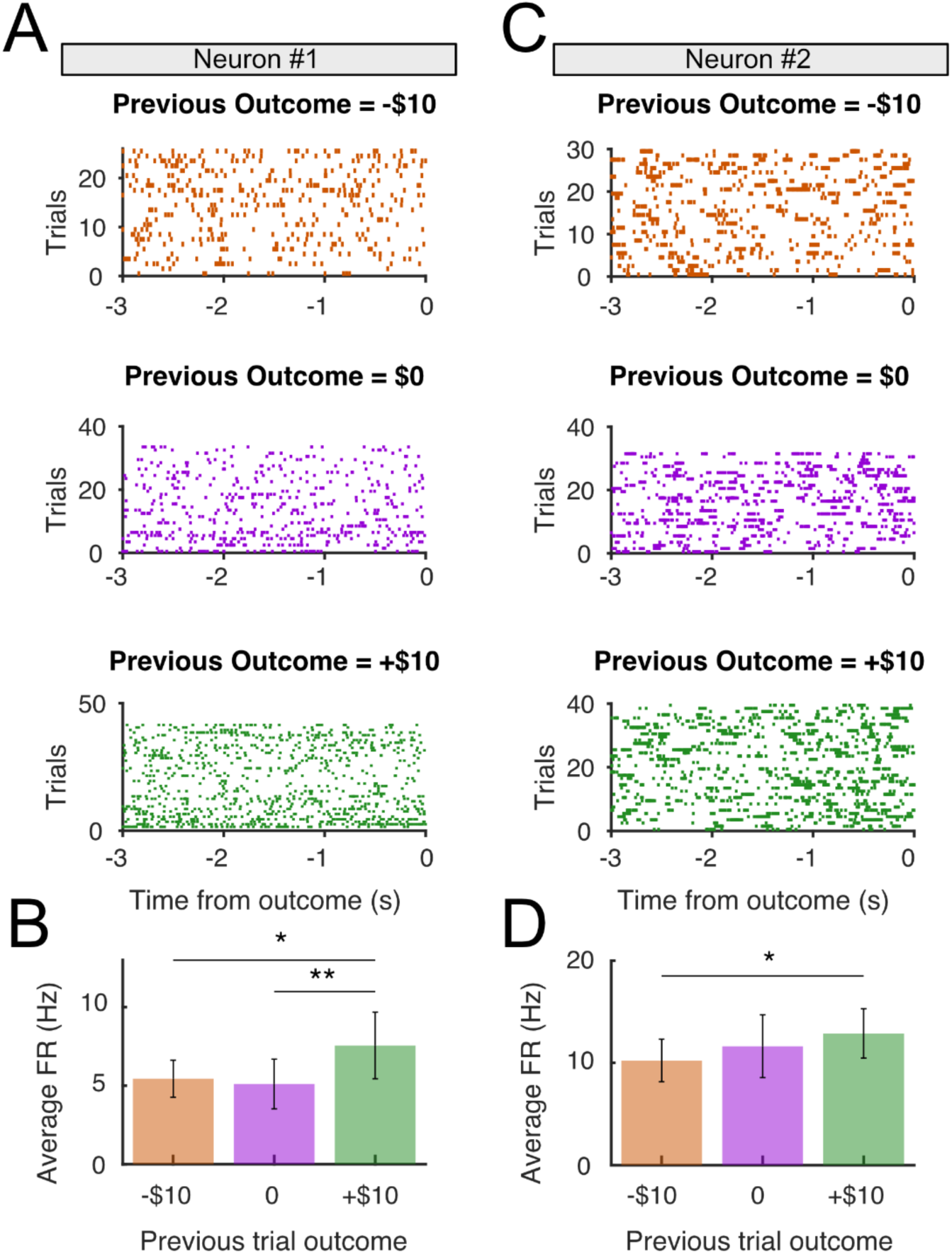
Sustained firing rate during the expectation period is modulated by previous reward outcome. (**A**) Activity raster plots from a representative neuron, separated according to previous trial outcome: negative (-$10, top), neutral ($0, middle) and positive (+$10), bottom during the expectation period prior between patient choice (t=-3000ms) and reward reveal outcome (t=0). (**B**) Firing rate during the expectation period split by previous reward outcome, averaged across trials. The sustained firing rate was significantly greater in trials following positive outcome than in trials following negative outcomes (p < 0.05, paired t-test) and neutral outcomes (p<0.01, paired t-test). **(C-D)** as **(A-B)**, for a second neuron that only showed differences between negative (-$10) and positive (+$10) previous outcomes.

Previous studies in rodents^8,27–29^ and humans^6^ have demonstrated that individual DA neurons reflect unexpected reward outcomes through FR modulation. To verify whether this was the case in our dataset, we also examined putative DA responses after reward outcome (feedback epoch) and compared FR after rewarded (i.e., positive outcome trials) vs unrewarded (i.e. negative outcome trials). In this feedback epoch, we found a subset of SN neurons (7/22, or 31.8%) whose FR was modulated according to the reward outcome (Supplemental Figure 2). Interestingly, whereas some showed an increase in FR following unexpected rewards (4/22, or 18.2%), a different set of neurons showed the opposite pattern, with FR decreasing following rewarded trials (3/22, 13.6%). We also conducted a GLM analysis similar to the one carried out during the expectation period at the post-reward outcome epoch. We included as regressors the current action (left or right), current trial reward identity (win or loss), current trial reward outcome (+10, 0, or - 10), and current RPE on the FR during the feedback period, none of which had a significant effect on FR during the feedback period (all p>0.05). Overall, we found limited evidence for reward encoding at the population level, which was especially salient in the mixed block where the value difference between the different potential outcomes was maximal (Supplementary Figure 3). Therefore, we found some evidence for reward encoding in putative DA neurons but with a certain amount of variation at the individual neuronal response level.

### The population level encoding of reward expectation and invigorating effect of higher DA signal

Next, we examined whether the expectation encoding was present at the population level by comparing the FR of all putative DA neurons during the expectation period across previous outcomes identity (-$10/$0/+$10). To compensate for the heterogeneity in baseline FRs across neurons, we normalized FRs to the average expectation period FR per neuron, across all trials. We found that FRs of putative DA neurons was higher in trials following positive (+$10) vs negative (-$10; p<0.01, one-tailed paired t-test) and neutral outcomes ($0; p<0.05, one-tailed paired t-test; Figure 4A-B). There was no significant difference between trials following negative and neutral outcomes (-$10 and $0, p=0.26, one-tailed paired t-test). Because some previous studies have shown a pre-reward ramping up of DA activity as animals get progressively closer to reward^17,30–35^, we also examined whether ramping dynamics existed in the FRs of our SN neurons by carrying out a linear regression between FR and time elapsed during expectation, in which a significant positive relationship would indicate the presence of ramping dynamics. However, we failed to see such an association (p=0.445), indicating that ramping dynamics were not clearly present. Finally, because previous evidence and numerous models of DA function relate tonic DA levels with response vigor^36^, we examined whether differences in prior outcomes also affected reaction times (RTs). We found that RTs were faster following trials with positive outcomes compared to after neutral/negative outcomes (+$10 median = 0.916s vs -$10/$0 median = 1.017s, p<0.05, Wilcoxon rank sum test, Figure 4C).

**Figure 4.**
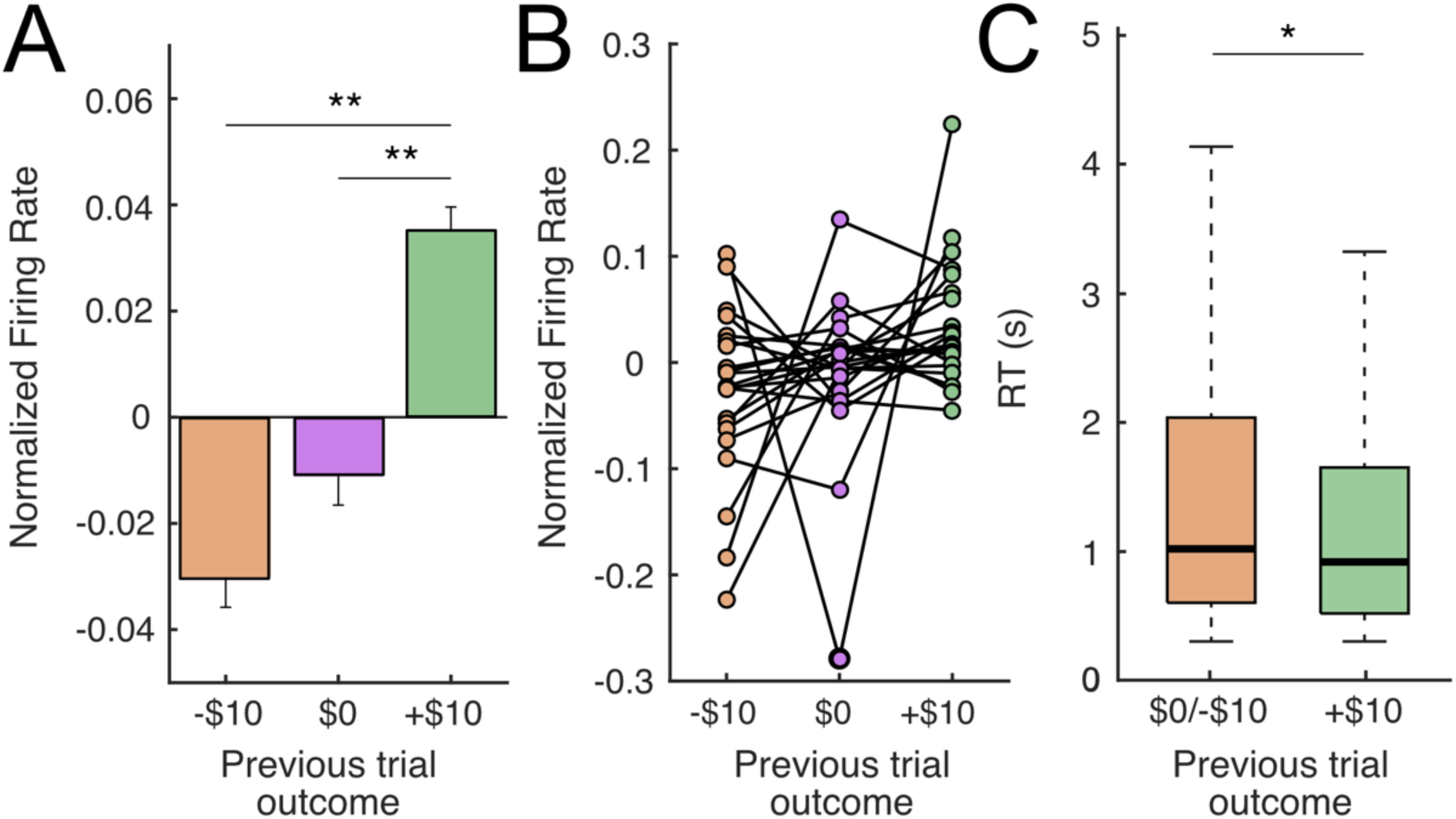
Putative DA neurons encode previous reward outcome during the expectation period. **(A)** Average normalized firing rate across all putative DA neurons during the reward expectation period (3s post-choice, prior to reward outcome reveal), separated according to previous trial outcome: negative (-$10), neutral ($0) or positive (+$10). FR of putative DA neurons is higher for previous positive outcomes than either previous neutral (p<0.05, paired t-test) or previous negative outcomes (p<0.01, paired t-test). **(B)** Same data as **(A)** but showing individual putative DA neurons. **(C)** Patient reaction times (RTs) varied depending on the previous trial outcome. RTs were faster following a positive, compared to neutral or negative outcomes (+$10 median = 0.916s vs -$10/$0 median = 1.017s, p<0.05, Wilcoxon rank sum).

Together, these results demonstrate that the sustained FR of SN neurons is modulated during the reward expectation period, with higher FRs during the expectation period following positive outcomes. Besides this neural effect, previous reward outcomes also had a behavioral correlate, with faster responses following positive outcomes.

## Discussion

The ability to compute differences between expected and obtained rewards is fundamental for adaptive decision-making in uncertain environments. This computation critically depends on ongoing, adaptive expectations of reward associated with risky choices. Despite this, the neural representation of reward expectations in the human brain remains elusive given the difficulty of recording electrophysiological activity from subcortical regions centrally involved in prediction error signaling, most centrally the dopaminergic system. Here, we build on previous intracranial single unit recordings^6,7^ and fast-scan cyclic voltammetry approaches^19,20^ that have started to unveil the dopaminergic dynamics underlying human economic decision-making tasks by leveraging invasive neurosurgical interventions. Specifically, we set out to examine whether human SN neurons explicitly encode reward expectations by recording spiking activity from putative DA neurons during a risky decision-making task.

Our results show that patients followed behavioral strategies that were strongly dependent on the previous trial outcome, often repeating choices machines that were previously rewarded. This behavior was accompanied by a modulation of putative DA spiking during the reward expectation period, with higher FRs following rewarded vs neutral or unrewarded trials, consistent with explicit encoding of a reward expectation signal. Finally, this increased activity was accompanied by faster RTs after rewarded trials, indicating a parallel effect of rewards on putative DA neuronal activity and response vigor.

### Behavioral modeling shows previous reward outcome impacts choice

We used a computational modeling approach to understand the behavioral strategies employed by patients in the two-armed bandit task. The modeling results show that patients’ behavior could be adequately captured by a win-stay/lose-shift heuristic in which their choices were primarily dependent on the reward outcome in the previous trial or by a Rescorla-Wagner (RW) model. The RW model fits had a high learning rate (mean = 0.73), meaning the immediately prior trial had a strong impact on their subsequent choices, with earlier trials being strongly discounted and, in many cases, not attended to (7/13 patients had a learning rate = 1). Therefore, although the quality of model fits to either of these models was very similar, meaning we could not definitively adjudicate patients’ behavior to either one, the behavioral predictions of the two models were similar. Specifically, under either of these behavioral strategies current choices are strongly dependent on the preceding outcome, with patients most often choosing a rewarded machine. Conceptually, the previous trial outcome can be thought of as a reward expectation signal, with previous positive outcomes leading to optimistic expectations and a higher proportion of ‘stay’ choices, whereas negative outcomes lead to pessimistic expectations and strategic switches.

### Spiking activity of individual putative DA neurons reflects previous trial outcome

Next, we observed a neurophysiological correlate for the behavioral weight of previous trial outcomes. Specifically, our results show that the FRs of putative DA neurons was modulated by the reward outcome of the preceding trial: FRs increased following positive outcomes (+$10) compared to neutral ($0) and negative (-$10) outcomes (Figure 3). Therefore, previous trial outcomes had a lingering effect on the subsequent activity of nigra neurons during the following expectation epoch. This effect was dependent on the sign of the reward outcome, with positive/negative outcomes leading to higher/lower FRs during the subsequent expectation period, consistent with the hypothesis that reward expectations derived from recent reward history are encoded in the sustained FR during the expectation period. These results are consistent with observations in animal models that tonic firing of DA neurons track values, with sustained firing increases following signaled value increases^17,32,37^ and tonic activity encoding the net reward rate.^36^ Recent evidence from animal models shows that DA transients in the striatum track reward rates, but potentially not in VTA DA responses^32^, whereas our results in contrast suggest that human SN neurons can signal reward expectation rates. In addition to value encoding, this tonic activity may influence motivational state and facilitate motor responses in high-reward context following positive outcomes, consistent with the proposed role of dopaminergic signaling in reward-related vigor.^38^ Consistent with this idea, we found that patient reaction times were significantly faster following positive outcomes (Fig. 4C), and therefore associated with increased FRs of putative DA neurons.

Overall, our results support the notion that increased tonic dopaminergic rates increased vigor and resulted in faster reaction times and provide the first demonstration of this effect at single neuron level in humans. In a more general sense, FR modulation during expectation can be related to observations that sustained neural activity can represent behaviorally relevant context, an effect that’s been observed in primate amygdala neurons^39^ and in high-frequency activity in human orbitofrontal cortex^40^.

### Heterogeneity in reward responses in putative DA neurons

Our study focuses on the pre-reward period, whereas most human electrophysiology studies have focused on the post-reward period. While we observed responses resembling reward prediction errors in some dopaminergic neurons (Supplemental Figure 2-3), post-reward responses were less robust than previous findings^6^, with a minority of neurons (∼30%) showing post-reward responses, and no statistically significant population-level reward responses. There are several potential explanations for this; first, our behavioral task was more complex than those previously used, with multiple block conditions. Therefore, it is possible that there are within-block or contextual expectation effects that obscure RPE encoding. Similarly, the block design reduced the number of trials available within condition (35 per block) and our ability to demonstrate statistically significant RPE encoding. However, these differences were qualitative rather than quantitative, as we did observe reward encoding, both at the population level (Supplemental Figure 3) and at the individual neuron level, with neurons showing phasic responses to the outcome (Supplemental Figure 2). Interestingly, we observed heterogeneity in the sign of these responses, with some neurons (n=3) showing a phasic increase in activity in response to unexpected reward and a similar number (n=4) showing a decrease. This is consistent with animal reports that indicate that there is significant heterogeneity in the encoding properties of dopaminergic neurons, with individual neurons capable of reflecting positive, negative RPEs or unsigned salience^41–45^, although there are likely to exist differences in the representation of RPE in SN and VTA^46^. Future experiments focused on characterizing the heterogeneity in response of putative DA post-outcome responses in the human SN, will constitute a promising future direction. Finally, we identified a large proportion of putative DA (n=22/27) vs putative GABAergic neurons (n=5/27) neurons in our sample, suggesting that our intraoperative selection criteria for individual cells (see Methods) successfully biased our recordings towards cells with lower FRs, which would fall primarily under a DA categorization under our criteria.

### Integration of expectation and reward responses in putative DA neurons

According to simple RPE models, RPEs arise from the arithmetic combination of expectation and reward information^1,47^ but the nature of their neural representations remains to be fully determined. Some models assume purely independent representation of both quantities converging at DA neurons^1,47–50^. Alternatively, reward and reward expectations may be computed redundantly in several different regions^16^, with other expectation and reward signals combined in upstream input regions^51–53^. Our data show that an expectation signal is explicitly encoded by the sustained FRs of putative DA neurons during the expectation period. Therefore, even though upstream regions show mixed selectivity of expectation and reward information in animal models, our data shows that human SN neurons can reflect individual components of the RPE calculation and are capable of explicitly encoding reward expectation signals.

### A proposed model of reward prediction error calculation

Despite these relatively sparse responses during the reward outcome period (Supplementary Figure 4), our observations make a few testable predictions about the nature of post-reward encoding in dopaminergic neurons. At face value, our results suggest that the reward response, present as a variable sustained FR during the expectation period, would not be simply ‘riding’ on top of the modulated expectation signal. Instead, the transient post-reward response should reach the same FR ceiling regardless of expectation. In other words, a similarly sized reward will always result in a similar maximum FR post-reward outcome, but the presence of different expectation period FRs during expectation would result in an adequately sized RPE signal. This is consistent with the model proposed in Hamid et al.^17^, whereby RPEs arise through dynamic modulation of baseline, not peak, DA signaling. Future invasive studies in human patients will be necessary to verify whether graded, expectation period-independent responses exist in human SN neurons.

In summary, by combining intraoperative SUA recordings and computational modeling of reward behavior, we provide evidence that human SN neurons explicitly encode reward expectations through sustained FR modulation, shedding light on the nature the neural representation of reward expectation in the human brain. Additionally, we provide evidence for an invigorating effect of positive reward outcomes which accompany these increases in SN firing. These findings add to our understanding of reward learning and motivational vigor in the human brain, with potential consequences for our understanding of psychiatric disorders in which the dopaminergic system is affected.

## Methods

### Deep Brain Stimulation Surgery

Research protocol was administered in accordance with the Icahn School of Medicine at Mount Sinai Institutional Review Board. We used intraoperative microelectrode recordings from patients undergoing DBS surgery for Parkinson’s disease. Patients were consented for research prior to their surgery date. Patients were off their medications at least 9 hours prior to the surgery as per surgical protocol. The instructions of the task were explained during the consent and again before playing the task. The surgery proceeded as per clinical routine and research was conducted when the patients were woken up for clinical testing during DBS targeting. A tungsten microelectrode with power-assisted Microdrive was lowered through the brain following the tractography previously determined by the clinical team.^54^ The surgeon (B.K.) lowered the electrode into the SNr after determining the ventral border of the STN and then listened for unit firing to confirm the location of the electrode (**Figure 1A**). The recordings were done from the dorsal substantia nigra. Once the surgeon was satisfied that stable spikes could be detected, the electrode was stationary for the duration of the experiment. To maximize our chances of recording from dopaminergic neurons which have lower FRs, we actively sought quieter neurons with lower spike frequencies, versus highly active neurons which were likely to be identified as putative GABAergic neurons. The patient was woken up during the surgery once the electrode was implanted for clinical testing, at which point they performed the task. Research time during the surgery was limited to 15 minutes or less (average = 10.30±2.48 min).

### Behavioral Paradigm

Patients played a two-armed bandit task in which they had to press a button to choose between two slot machines with initially unknown reward probabilities. Patients were taught the layout of the task and had a chance to practice in the days prior to their surgery and then again right before starting the game during surgery. Patients were instructed to maximize their reward by finding and choosing the machine that had the higher probability of resulting in the better outcome. The task had three blocks with 35 trials per block: a reward block, a punishment block, and a mixed block, whose order was randomized across patients. Each block had different win and loss outcomes; the reward block had positive (+$10) or neutral (+$0) outcomes; the punishment block had negative (-$10) and neutral (-$0) outcomes, and the mixed block had positive (+$10) and negative (-$10) outcomes. The two machines had different probabilities of resulting in the better outcome. One had a high probability (80%/20%) and the other had a low probability (20%/80%) (Figure 1B). These probabilities switched twice throughout each block on trials 12/13 and trials 24/25 depending on the reversal condition. The patients were made aware of this during the instructions to the task. Patients were taught the layout of the task in the days prior to their surgery and then again right before starting the game during the surgery. Patients were told the objective of the game was to maximize their reward by finding and choosing the machine that had the higher probability of giving the winning amount. The task took approximately 10 minutes on average (mean = 10.30±2.48 minutes), and patients were not compensated for participating in research as per IRB protocol. For reaction times analyses, we measured the time between trial presentation and choice and excluded trials with response times shorter than 300 milliseconds and longer than 12 seconds which are likely due to patient error.

### Computational Modeling of Behavior

For each patient, we first calculated model-free metrics of behavior: number of wins and losses in each block as well as the number of times the machine with the higher reward probability was picked regardless of whether it actually resulted in the better outcome.

To determine which learning method our patients were using to make their decisions we carried out model comparison. We evaluated a random model in which patients chose randomly between the two options, a heuristic win/stay-lose/shift model, and a Rescorla-Wagner model. We first validated the models testing them with simulated data for each model based on the parameters of the task. For the random model, actions were selected randomly whereas with the heuristic model actions were simulated based on the outcome produced by the previous action (either a win or loss). In the Rescorla-Wagner model, the agent is expected to use the outcomes from previous trials to determine the values of each slot machine and act according to those expected values. The parameters examined in the Rescorla-Wagner model are the learning rate (⍺) and the inverse temperature (β) which signifies the level of predictability of choice^55^. The value update function can be written as:

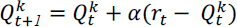

where α indicates the learning rate, which captures the extent to which the prediction error, updates the cached value, ranging from α = *0* for a non-learner agent to α=1for maximal learning from the prediction error. The RPE is captured by the term 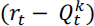, indicating the difference between the expected 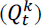 and the obtained (*r_i_*) reward. Therefore, the higher the learning rate, the faster an agent updates their values and takes more recent outcomes into account, whereas a lower learning rate means the agent updates their values at a slower pace and looks at the trend of outcomes beyond the most recent one. The mapping of values to choices is achieved through the following SoftMax equation:

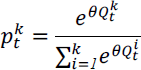

where θ is the ‘inverse temperature’ parameter that controls the level of stochasticity in the choice, ranging from θ = *0* for random responding to θ = *∞* for deterministically choosing the highest value option. The learning rate is dependent on both the unconditional and conditional stimuli.^25^

Following data simulation, we did parameter recovery for the Rescorla-Wagner model. Parameter recovery involves using the simulated data with known parameters and fitting the model with the simulated data to recover the parameters back again. The recovered parameters are then compared to the initial parameters to check for correlation. If there is a high correlation, that confirms that the model parameters are identifiable from the task^56^. Finally, we compared the models using a confusion matrix which allows us to see if the data simulated using a certain model is best fit only by that model. If data is better fit by another model that could mean that model is not identifiable with the data and the experimental parameters^56^. We followed these steps first for the entire task without splitting by block followed by splitting by block. We used the outcome to these results to first check if the task was suited for a RL model or if a more complex learning model would be required to model this task. Secondly, by estimating the best model for each of our patients’ behaviors, we can confirm that our patients understood the objective of the task and were making informed decisions as opposed to selecting randomly.

### Electrophysiological Data Collection

Neural data was recorded from an Alpha Omega machine (Alpha Omega, Nazareth Illit, Israel) routinely used for microelectrode recordings in our DBS cases. Neural data was digitized at a rate of 44000 Hz and filtered between 300-9000 Hz to isolate high-frequency activity reflecting neuronal spiking. Filtered data was analyzed with the OSort spike sorting software^57^. Briefly, spikes were detected if the bandpass filtered signal crossed a threshold determined as five times the standard deviation of the average amplitude of the signal. Once a new spike was detected, the multidimensional distance from the spike to clusters of similar spikes already formed was calculated. If the distance was smaller than a threshold calculated as the square of average standard deviation of the filtered signal calculated along a sliding window times the number of datapoints in a single waveform, then it was added to an existing cluster, but if it was larger than the threshold a new cluster was formed^57^. The clusters were then manually examined for artifactual waveforms, or clusters that were artificially split and merged if necessary (Figure 1C). Clusters were also determined to be artifactual if their interspike interval was within 3 ms greater than 5% of the time. To determine if clusters were artificially split, we analyzed the projection tests outputted by OSort. The projection test compares each cluster against each other by looking at the distance between each cluster and examining if the clusters are far apart enough to consider them two separate clusters.

To distinguish between dopamine neurons and GABAergic interneurons, we calculated the average FR of individual neurons. Putative DA neurons were classified as having an average FR less than 15 Hz, and putative GABAergic interneurons were classified as having an average FR greater than 15 Hz^22^. Neurons with an average FR less than 0.5 Hz were discarded.

The neural recordings were epoched time locked to the events in the game. We focused on a range of -3000 ms-1000 ms around outcome reveal. That time frame was chosen because the time in between choice and outcome reveal was 3000 ms and then the outcome stayed on the screen for 1000 ms. Raster plots showing timing of individual spikes were generated for each neuron per block. We then computed the z-scored FR for the average response curve pre, and post outcome reveal for win and loss trials per block. Z scoring was done by normalizing the FR against the average expectation period FR for the neuron.

## Supplemental Figures

**Supplemental Table 1.**
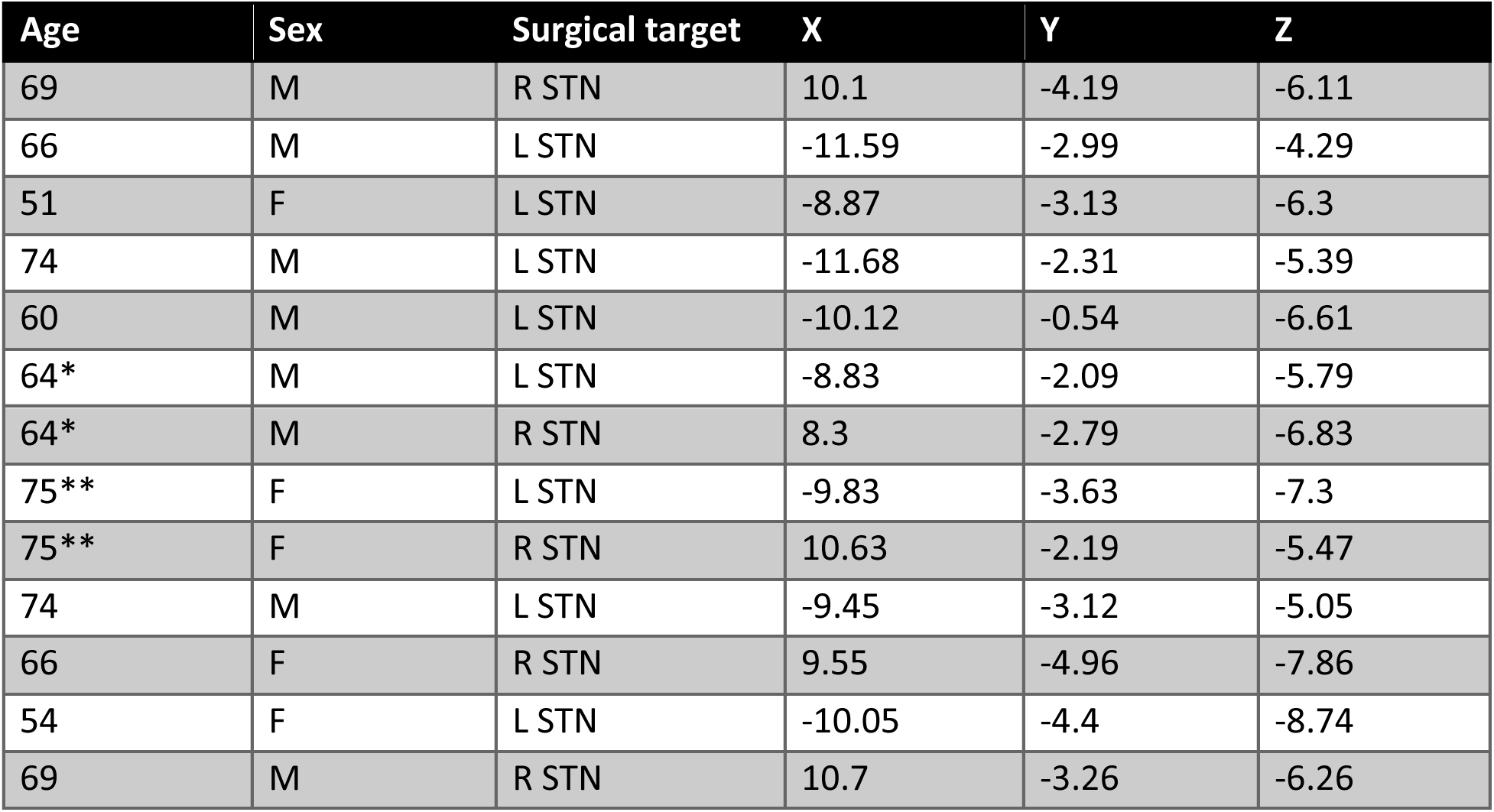
Patient statistics. Details for patients that participated in the study including age, sex, surgical target, and x, y, z coordinates for the microelectrode recordings relative to the middle cerebellar peduncle (MCP). * and ** represent different sides of the brain for the same patient.

**Supplemental Figure 1.**
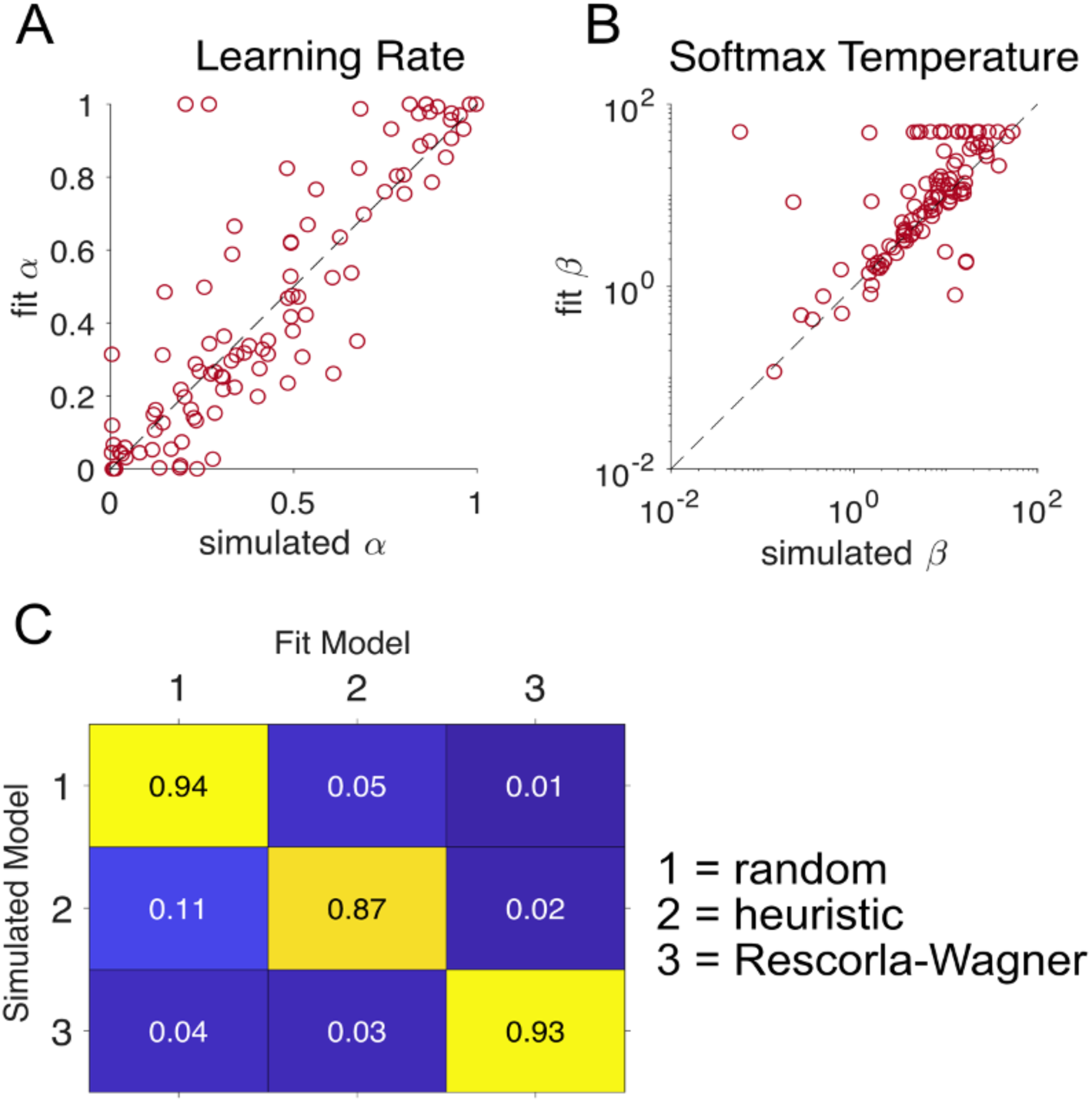
Computational modeling of task and RW parameter recovery. Parameter recovery showing correlation between the **(A)** fit and simulated learning rate and **(B)** temperature. **(C)** Confusion matrix shows that all 3 models are distinguishable from each other with the given task parameters. The RW model parameters are recoverable, and all 3 models are distinguishable from each other with the given task parameters.

**Supplemental Figure 2.**
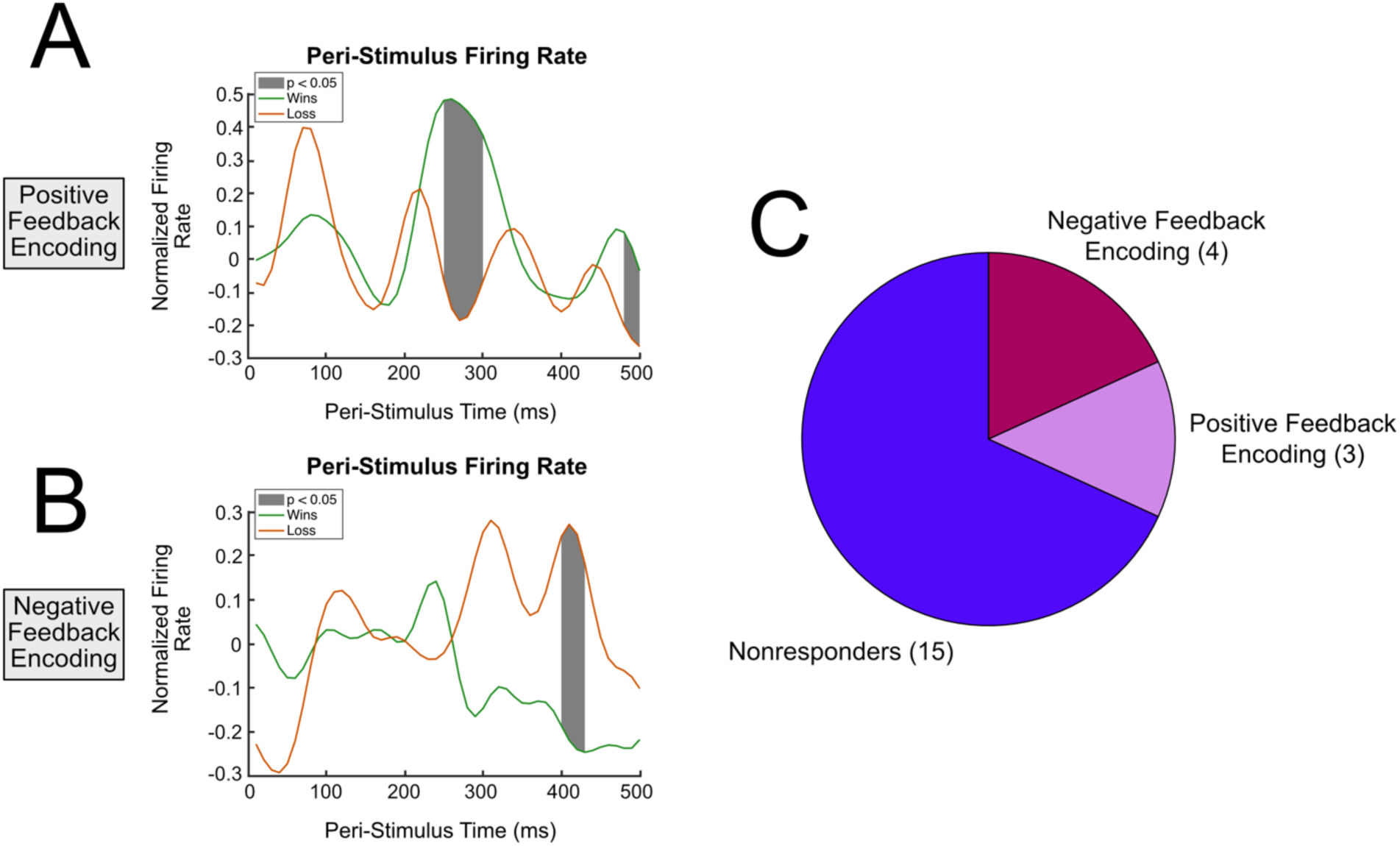
Breakdown of encoding properties for individual neurons shows heterogeneity in responses. **(A)** A positive feedback encoding neuron. **(B)** A negative feedback encoding neuron. **(C)** Distribution of individual neuron encoding properties. Majority of neurons were non responders.

**Supplemental Figure 3.**
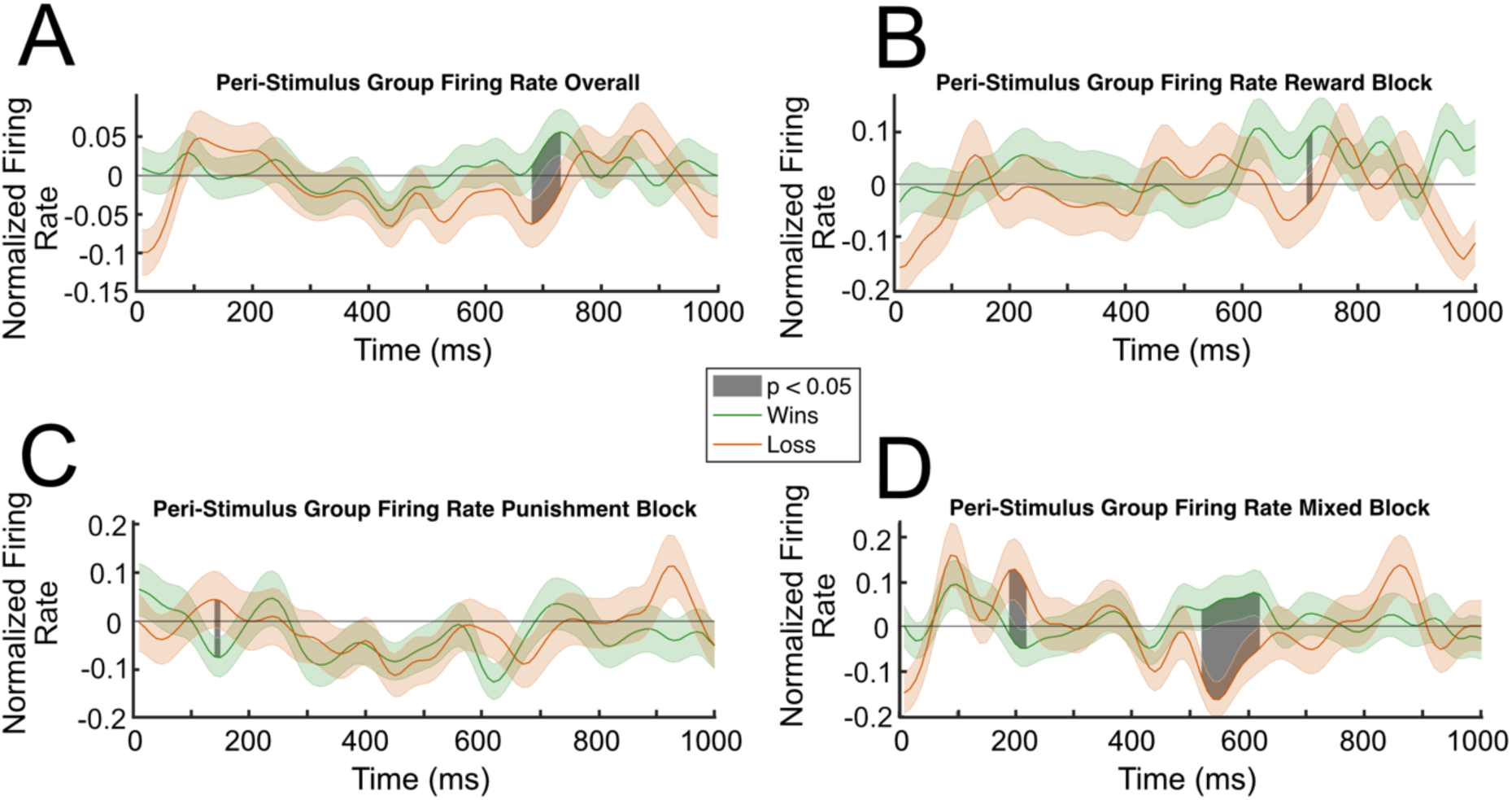
Win and loss trials show no consistent significant difference in firing rates. **(A)** Win and loss trials, collapsed across all blocks. Win and loss trials for **(B)** rewards block, **(C)** punishment block, and **(D)** mixed block. Significance was found using a permutation test.

**Supplemental Figure 4.**
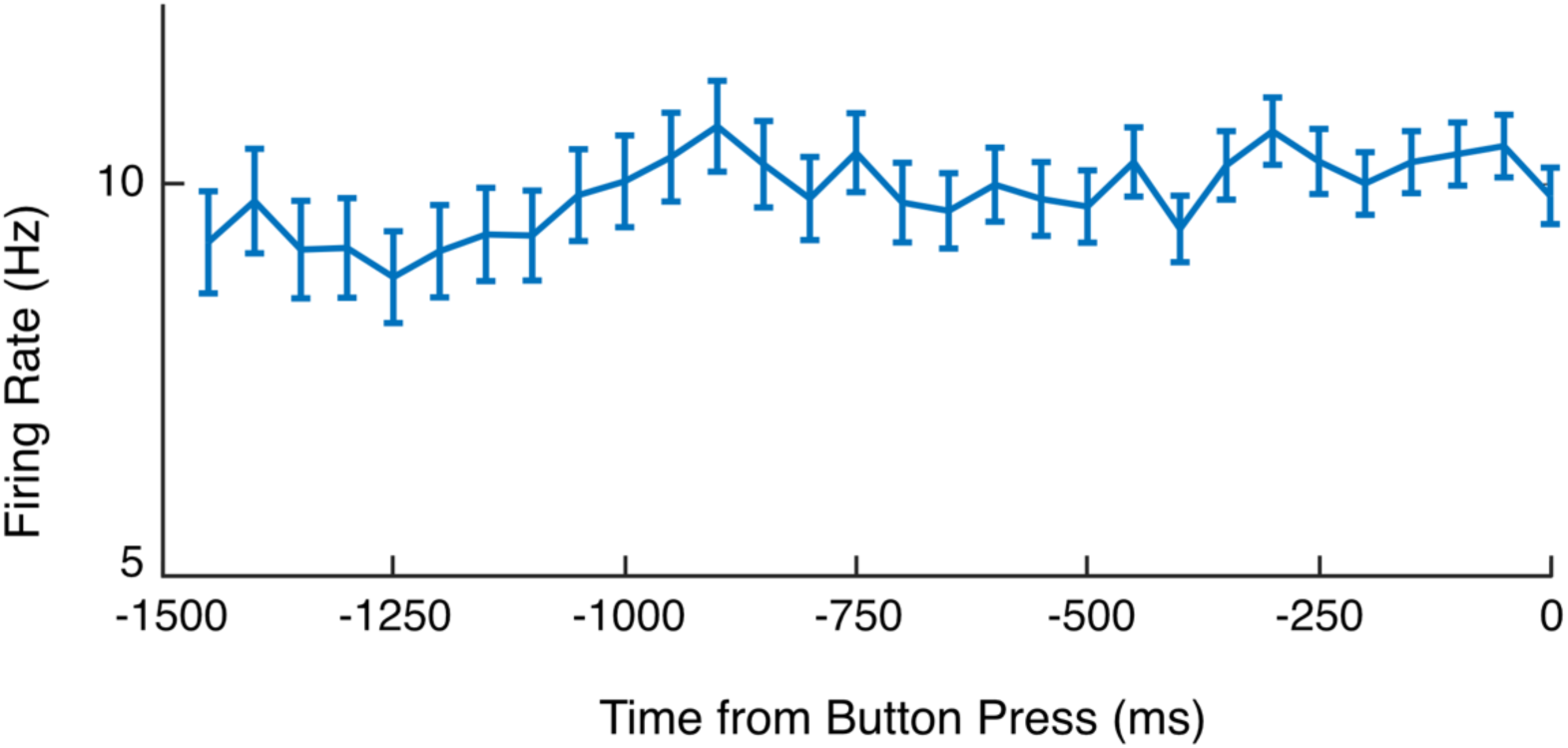
Average firing rate across all trials leading up to button press shows no movement related activity. The average firing rate in the 1.5s leading up to the button press. There is no increase or ramping effect of neuronal activity related to movement as the participant gets ready to press the button.

## Acknowledgements

We are grateful to all patients that participated in this study. We would like to thank Juri Minxha for assisting in the use of OSort and the surgical team at Mount Sinai West for support during data collection. We would also like to thank Alice Hashemi and Kimia Ziafat for consenting the patients for the research. We are grateful to Naoshige Uchida and members of the Saez and Gu laboratory for their comments on the manuscript. A.K. is supported by JSPS Overseas Research Fellowships. X.G. is supported by the National Institute on Drug Abuse [grant numbers: R01DA043695, R21DA049243] and National Institute of Mental Health [grant number: R21MH120789, R01MH122611, R01MH123069, R01MH124115]. I.S. is supported by the National Institute of Mental Health [grant numbers: R01MH124763, R01MH104606] and the National Science Foundation [project number: 2319580].

## Author Contributions

I.S, X.G, and B.H.K designed the study. Z.I., L.N.M, A.N.D, M.H., A.M., and B.H.K. were involved in data collection. K.K. and X.G. designed the task. Z.I., S.E.Q., A.K. and A.N.D. carried out neural data analyses, Z.I., S.E.Q. and A.K. carried out behavioral analyses. I.S. and X.G. supervised all analyses. B.H.K. supervised surgical aspects of the project. Z.I., A.K., and I.S. wrote original draft of the paper. All authors participated in editing and approved the final document.

